# African swine fever virus I196L is a virulence determinant and its deletant induces robust protection in Domestic pig

**DOI:** 10.1101/2023.06.22.546121

**Authors:** Jiaqi Fan, Rongnian Zhu, Nan Li, Jinjin Yang, Huixian Yue, Yanyan Zhang, Xintao Zhou, Junnan Ke, Yu Wang, Qixuan Li, Yu Qi, Faming Miao, Min Li, Teng Chen, Rongliang Hu

**Author notes:** Correspondence (R.H.). Jiaqi Fan, Rongnian Zhu and Nan Li contributed equally to this work.

## Abstract

The worldwide pandemic of African swine fever virus (ASFV) has a profound impact on the global pig industry. ASFV is a complex multilayered structure and the functions of unknown genes are being revealed. Here we deleted I196L from virulent ASFV SY18 with different length and obtained two recombinant viruses. The replication efficiency of the two recombinant viruses were similar but significantly lower than parental SY18. The pigs all survived the two recombinant viruses with 10^6.0^TCID_50_ except one pig occurred sudden death and the suvived pigs all resisted the challenge without fever after intramuscularly injecting a lethal dose (10^2.0^TCID_50_) of ASFV SY18. The recombinant viruses induced a strong anti-p54 humoral immune response. Meanwhile, the pigs also inevitably appeared moderate to high viremia throughout the observation period and presented a gradually downward trend. The results show that deleting I196L gene is a potential and effective vaccine that protects pigs from ASFV.

**IMPORTANCE:** The worldwide outbreak of African swine fever (ASF) cannot be effectively prevented due to no availably commercial vaccine. Many different types of vaccine candidates are researched and reported, which is a hopeful trend to develop safety and efficacy vaccine. Here we report on an unknown functional gene, I196L, which affects the virulence and replication of ASFV. When I196L was deleted from ASFV SY18, the recombinant virus decreased virulence and resisted the challenge of parental strain. This is a novel, effective, potential live attenuated vaccine (LAVs) for ASF.

## INTRODUCTION

African swine fever (ASF) is high contact infectious, febrile and fatal disease caused by African swine fever virus (ASFV). ASFV is a structurally complex and enveloped double-stranded DNA virus with a genome between 170 and 194 kbp, which is the sole member of the Asfarviridae family, genus Asfivirus(1-3). ASFV has similar structure and replication strategy with poxviridae and iridoviridae(4). Tick is known carrier of the virus. The wild boar and domestic pigs are terminal host and domestic pig usually shows obvious clinical signs(5-7).

The majority of ASFV genes are unknown and have little homology with most known viral proteins. The B646L gene, encodes the p72 protein (ASFV structural protein), is commonly used for genotyping ASFV due to high homology. Currently, ASFV isolates are classified into 24 genotypes, 22 of which are endemic only in Africa(8). ASFV genotype I spread outwards from Africa to Europe and South America in the 1990s and was subsequently largely eliminated (except in Africa and Sardinia, Italy) through aggressive decontamination efforts(9, 10). ASFV genotype II spread outwards across the continent to the Caucasus in 2007, and then continued to spread uncontrollably in Europe, Asia, north Americas and Oceania(11-15). ASF was first confirmed in China in August 2018 and a number of prevalent highly pathogenic ASFV strains are monitored in recent years and show high nucleotide similarity.

There is still no acceptably commercial vaccine for ASF at present. The quarantine and trapping are still used to stop the spread of ASF. Several natural and genetically engineered attenuated strains have been reported and pig can obtain long-term protection against challenge by homologous virulent strain. ASFV-G-ΔI177L vaccine has been applied to market in Vietnam. It is necessary to develop effective live attenuated vaccines (LAVs)(16-18). Certainly, the safety of vaccine needs continued attention.

The deletion of a single key gene in ASFV genome can alter viral pathogenesis. The candidate strains of LAVs deleting single gene, such as ASFV-G-ΔI177L(19), ASFV-G-ΔA137R(16), SY18ΔI226R(20) and ASFV-GZΔI73R(21). Here, we report that deletion of a previously undescribed gene, I196L, from a highly virulent ASFV SY18 strain resulted in attenuation of virulence. The I196L gene is a conserved gene at the 3’ end of the ASFV genome and there is no relevant report on I196L natural deletion in known ASFV strains. We obtained the recombinant viruses SY18ΔI196L-1 and SY18ΔI196L-2 by replacing ASFV SY18 I196L gene with an enhanced green fluorescent protein (EGFP). The result shows that the SY18ΔI196L-1 and SY18ΔI196L-2 decreased the replication in primary alveolar macrophages (PAMs). More importantly, the two recombinant viruses greatly reduce the virulence and remains the immunogenicity.

## RESULTS

### Genetic diversity of I196L of ASFV strains

The I196L gene in ASFV SY18 consists of 609 bp and encodes 202 amino acid residues, which localized in antisense strand of the SY18 genome between 175047 and 175655 base pairs. To assess the genetic diversity of I196L, we analyzed the I196L amino acid residues of different genotypes of ASFV, which is between 69.68%-100% among the 16 ASFV genomes and the homology between the same genotype is high. The I196L consists of 202 amino acids with 100% identity in the ASFV genotype II and consists of 196 amino acids in the ASFV genotype I with 100% identity. I196L gene was highly conserved in genotype I and II. Among them, the amino acid sequence of I196L in ASFV genotype II was consistent with that of Esontia 2014 (Genbank: LS478113), Pig/HLJ/2018 (Genbank: MK333180), Georgia/2007/1 (Genbank: NC044959). The amino acid sequences of I196L of the six strains of ASFV genotype I including OURT88/3 (Genbank: NC_044957), NHV (Genbank: NC_044943), L60 (Genbank: NC_044941), E75 (Genbank: NC_044958), Benin_97-1 (Genbank: NC_044956), BA71V (Genbank: U18466) were identical

### I196L is a late-stage transcriptional gene

The transcriptional kinetics of I196L mRNA were examined after infection. PAMs were infected with ASFV SY18 (MOI=3), and cell cultures were collected at 1, 4, 8, 12, 16, and 24 hours post infection (hpi), after which total RNA was extracted from the cells. Real-time quantitative reverse transcription PCR was used to detect the transcript level of I196L mRNA. The CP204L (p30), an early transcribed gene and B646L (p72), a late transcribed gene, were used as reference. The expression of the p30 gene reached the plateau at 8 hpi, and mRNA levels of I196L gene and p72 gene are still increasing at 8 h-16 h. The results suggest that I196L is a late transcribed gene in the ASFV genome. This result is consistent with the results of Cackett literature(22).

### Development of the recombinant virus SY18ΔI196L-1 and SY18ΔI196L-2

To investigate the function of the I196L gene, we designed two recombinant viruses, SY18ΔI196L-1 and SY18ΔI196L-2), lacking the I196L gene and replaced the I196L gene with EGFP. The I196L gene in ASFV SY18 strain has a total of 609 bp, and the I196L gene and I177L gene share 8 bp. The SY18ΔI196L-1 deleted 601 bp in the I196L gene (without affecting the ORF of I177L) (Fig. 3a), and the SY18ΔI196L-2 deleted all 609 bp of the I196L gene (Fig. 3b). The recombinant viruses were purified according to screening and diluting fluorescence-activated cells. The genomes of SY18ΔI196L-1, SY18ΔI196L-2 and parental strain were sequenced to verify whether the genomes change. The NGS results showed that the I196L gene in SY18ΔI196L-1 and SY18ΔI196L-2 had been successfully replaced as expected. No undesired modifications were detected in the SY18ΔI196L-1 genome. SY18ΔI196L-2 showed one mutation in the 379th base of E199L ORF resulting in the change in the 133rd amino acid from Gly to Arg.

**FIG 1.**
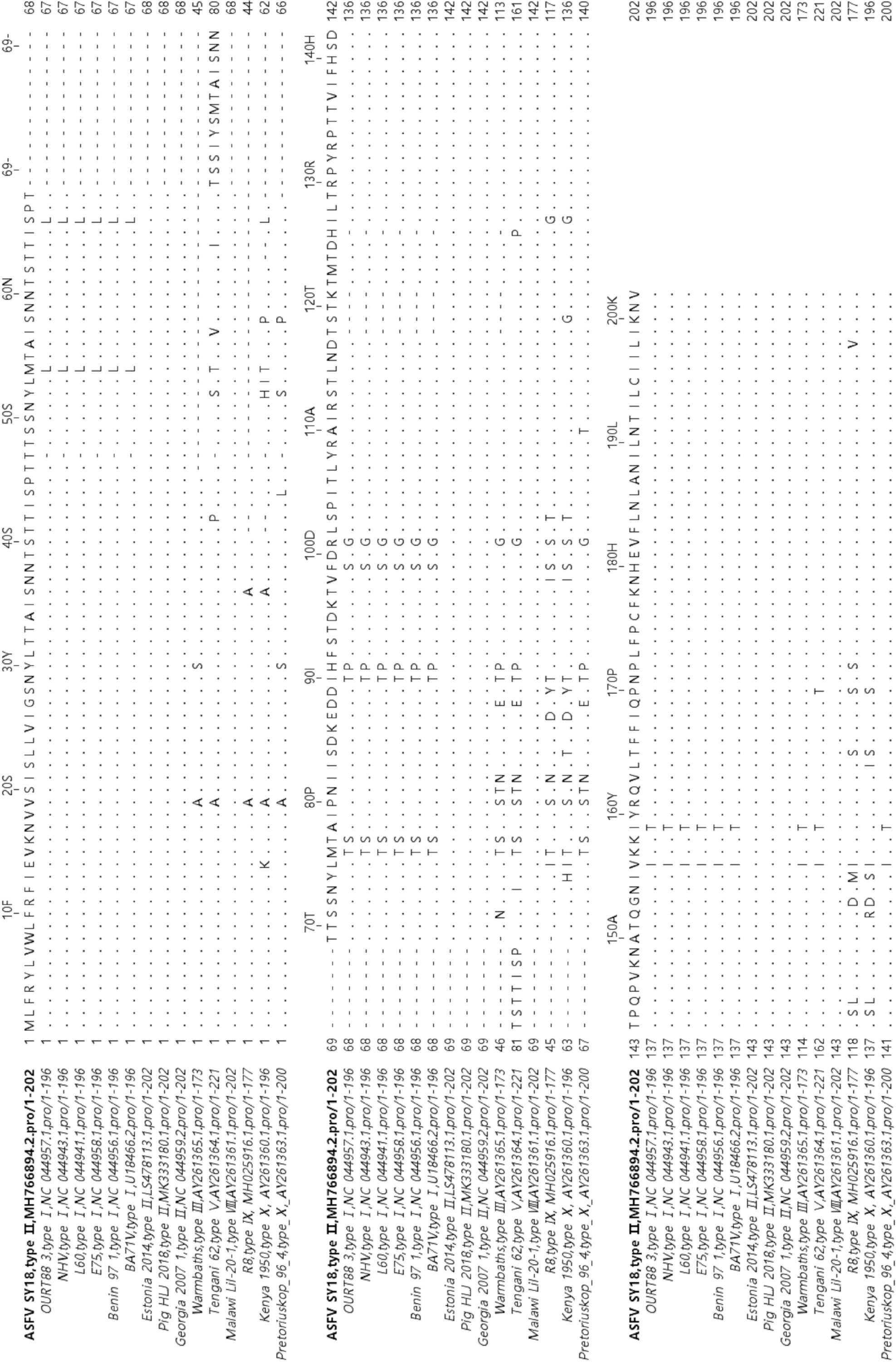
Multiple sequence comparison of I196L amino acid residues in 16 ASFV isolates. The amino acid sequence of I196L in ASFV SY18 was used as the reference sequence. Identical amino acids are indicated by “.”. Blank amino acids are indicated by “-”. Amino acid sequences were compared using Jalview software (accessed November 20, 2022).

**FIG 2.**
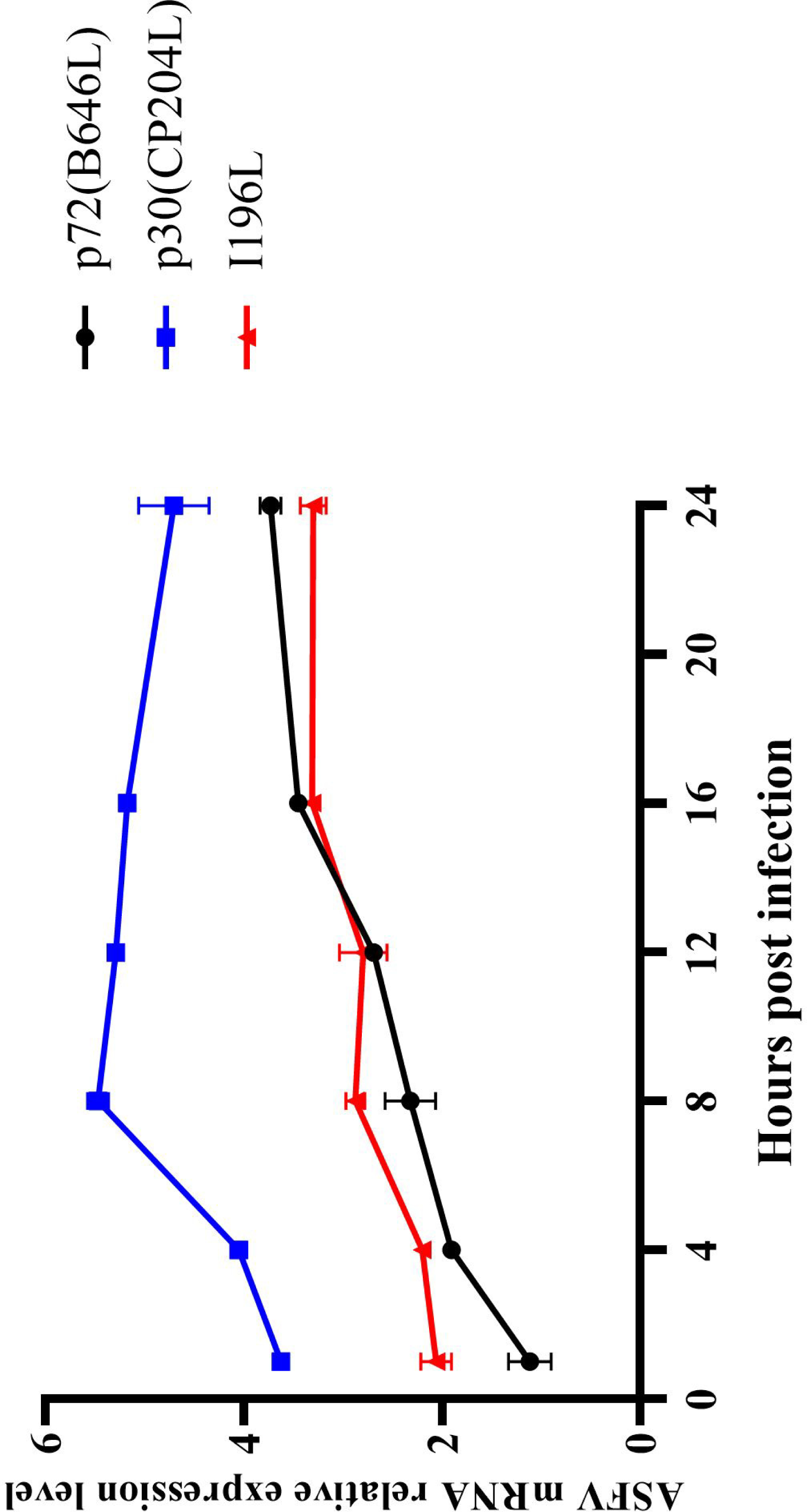
Transcript levels of the I196L gene during infection of PAMs by ASFV. The relative mRNA expression levels of I196L, CP204L, and B646L genes were detected by RT-qRCR at different infection times, meanwhile, the GAPDH gene was used as control.

**FIG 3.**
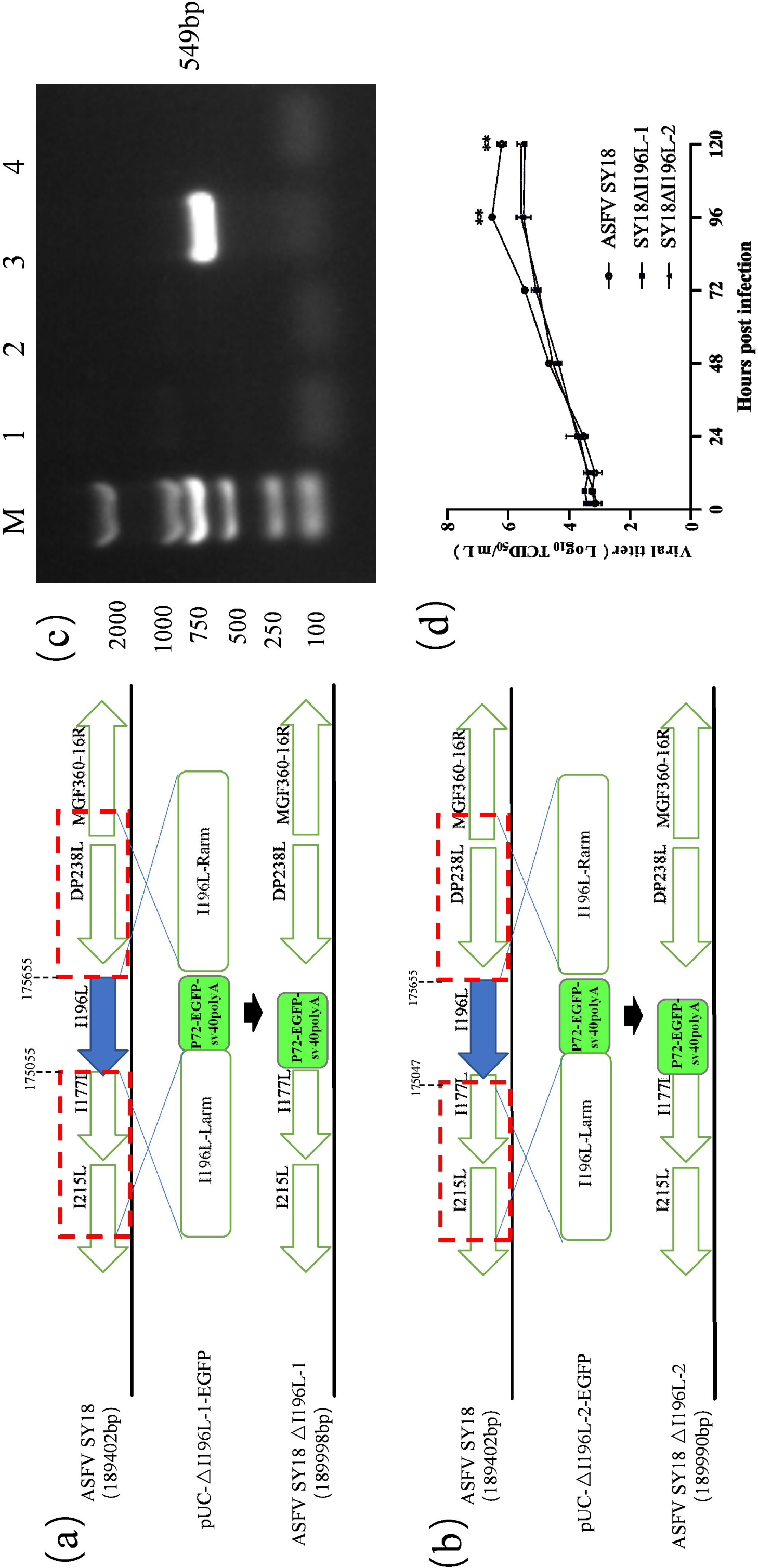
Construction and growth kinetic of SY18ΔI196L-1 and SY18ΔI196L-2. (a, b) Schematic diagram of the construction of SY18ΔI196L-1 and SY18ΔI196L-2. The PAMs were transfected recombinant transfer vector p72eGFPΔI196L and subsequently were infected ASFV SY18. The homologous recombination happened between p72eGFPΔI196L and ASFV SY18 genome and I196L gene was replaced with an EGFP cassette. (c) PCR amplification results of the I196L. The line 1, line 2, line 3 and line 4 were the PCR amplification targeting I196L gene of SY18ΔI196L-1, SY18ΔI196L-2, SY18 and no template control. (d) Growth curves of SY18ΔI196L-1, SY18ΔI196L-2 and parental virus. PAMs were infected with three viruses at 0.01 MOI and were determined virus titer at different infection time. The y-axis represented virus titer and exhibited with log10 TCID_50_/mL and the x-axis represented the hours post infection. The data are expressed as mean ± SD of 3 independent experiments. *0.01 < *P* < 0.05, \*\**P* < 0.01.

### Growth characteristic of recombinant virus in vitro

Two recombinant viruses and parental virus infected PAMs at 0.01 MOI and the samples were collected at 1, 6, 12, 24, 48, 72, 96 and 120 h and the viral titers in each sample were examined. The results showed that the parental virus reached plateau at 96 h about 10^6.5^TCID_50_/ml and the SY18ΔI196L-1 and SY18ΔI196L-2 were all about 10^6.5^TCID_50_/ml, which was significantly different from the parental strain (p < 0.05) (Fig. 3b). Therefore, though the I196L gene was not essential for ASFV replication, deleting I196L gene significantly reduced the replication of parental virus in vitro.

### The recombinant virus remains normal ASFV structure

Taking into consideration that the reduced replication, the ultrastructure of the recombinant viruses was examined by transmission electron microscopy (TEM). SY18ΔI196L-1, SY18ΔI196L-2 and ASFV SY18 infected PAMs were analyzed at 48 h. The results are shown in Fig. 4. The three viruses formed mature icosahedral structures in cytoplasm and the diameter of virus particles was about 200 nm (Fig. 4). There were a large number of viral particles of SY18ΔI196L-1 in PAMs (Fig. 4a, 4a-1 and 4a-2) and the “viral factory” area near the nucleus was clearly observed including abundant precursor virus membranes, immature viruses and mature viruses. Ribosomes are clearly observed around mature and immature viruses in the virus factory.

**FIG 4.**
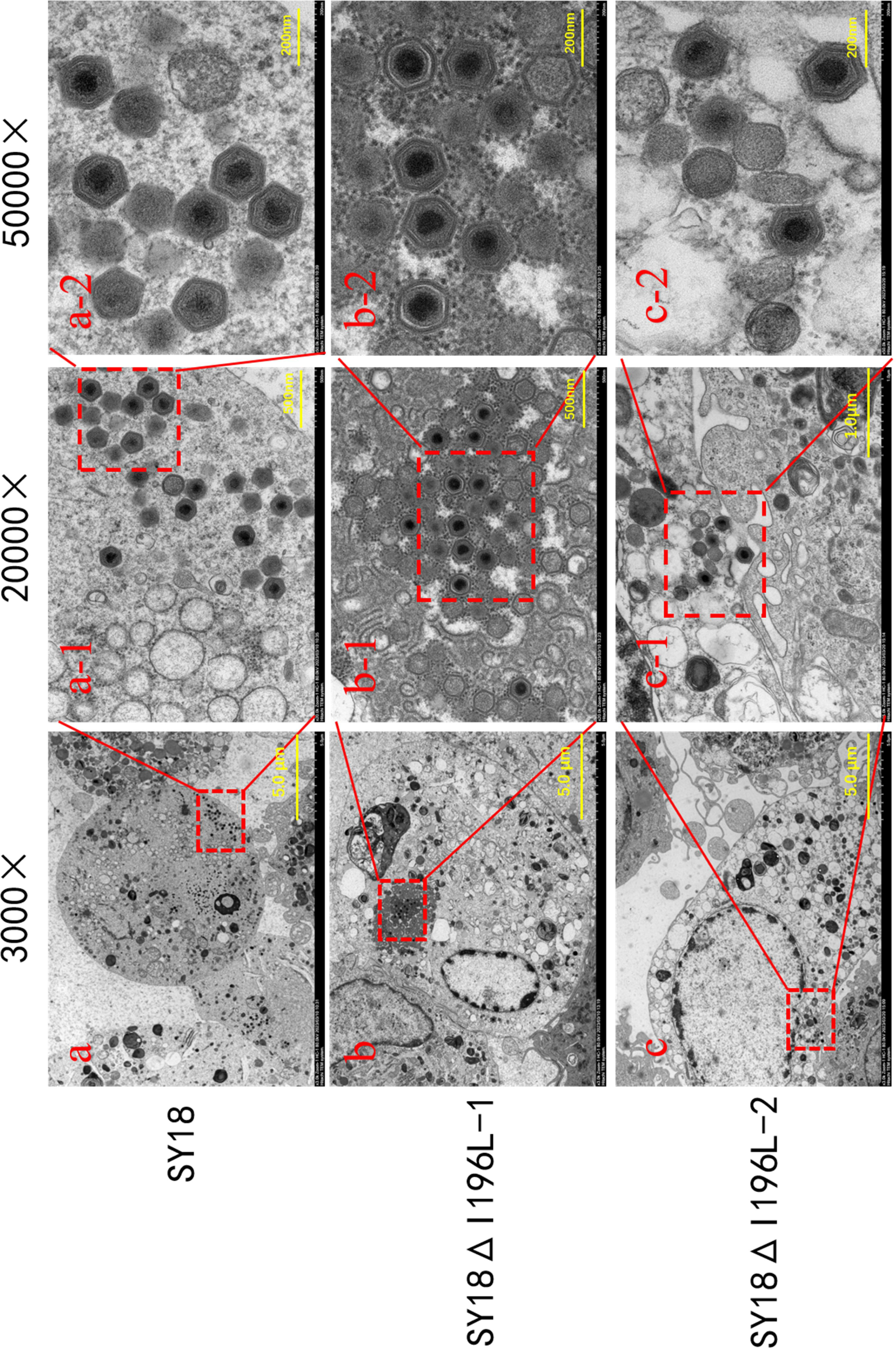
Ultrastructure of ASFV SY18, SY18ΔI196L-1 and SY18ΔI196L-2. (a, a-1, a-2) Ultrastructure of ASFV SY18 infected with PAMs for 48 h. The figures were progressively magnified by 3000×, 20,000× and 50,000× and showed a large amount of both immature and mature viruses. (b, b-1, b-2) Ultrastructure of SY18 ΔI196L-1. The magnified figures showed the immature and mature viruses at 3000×, 20,000× and 50,000×. (c, c-1, c-2) Ultrastructure of SY18 ΔI196L-2. The figures are magnifications at 3000×, 20,000× and 50,000× and showed the typical ASFV morphology of the different stages of infection.

### Evaluation of recombinant virus virulence in pigs and the efficacy of protection against parental ASFV SY18 challenge

Fifteen pigs in three groups were intramuscularly inoculated with 10^6^ TCID_50_ of SY18 ΔI196L-1 (group 1, G1), 10^6^ TCID_50_ of SY18 ΔI196L-2 (group 2, G2) and saline (group 3, G3). During the immunization, some pigs in G1 (4/5) and G2 (3/5) showed a transient fever starting on day 6 and lasting for 2 to 3 days. All pigs had no other symptoms associated with ASF during the immunization period (Table 1). However, pig G1-5 died on the 18th day of immunization owing to the stress caused by the stimulation of blood collection on the 14th day. The survival rate during immunization was 80% for group 1 and 100% for group 2. The deletion of the I196L gene caused a dramatic decrease in the virulence of the ASFV SY18.

**Table 1.** Clinical performance and survival rate after immunization

| Groups | No. of survivors/total | No. of fever/total | Mean value for fever parameter (SD)Fever ( $\geq 40.5^{\circ}\text{C}$ ) | | |
| --- | --- | --- | --- | --- | --- |
| | | | Day of onset (days±SD) | Duration no. of days (days±SD) | Highest rectal temp ($^{\circ}\text{C}\pm$SD) |
| $10^6$ TCID<sub>50</sub> SY18ΔI196L-1 | 4/5 | 4/5 | 6±1.22 | 2.25±0.43 <sup>a</sup> | 40.7±0.21 <sup>a</sup> |
| $10^6$ TCID<sub>50</sub> SY18ΔI196L-2 | 5/5 | 3/5 | 7.33±0.47 | 2±0.82 | 40.97±0.05 |

a : The pig G1-5 suddenly died due to artificial stimulation.

During challenge, pigs in G3 developed ASF clinical phenomena (fever, anorexia, diarrhea and skin bruising) on day 6 and died on day 11 (Fig. 5b). All pigs in G1 (4/5) and G2 (5/5) showed normal clinical signs (no fever and ASF-related signs) and survived challenge (Table 2). SY18 ΔI196L-1 and SY18 ΔI196L-2 dramatically reduced the virulence of ASFV SY18 and prevented homologous lethal challenge from ASFV SY18.

**FIG 5.**
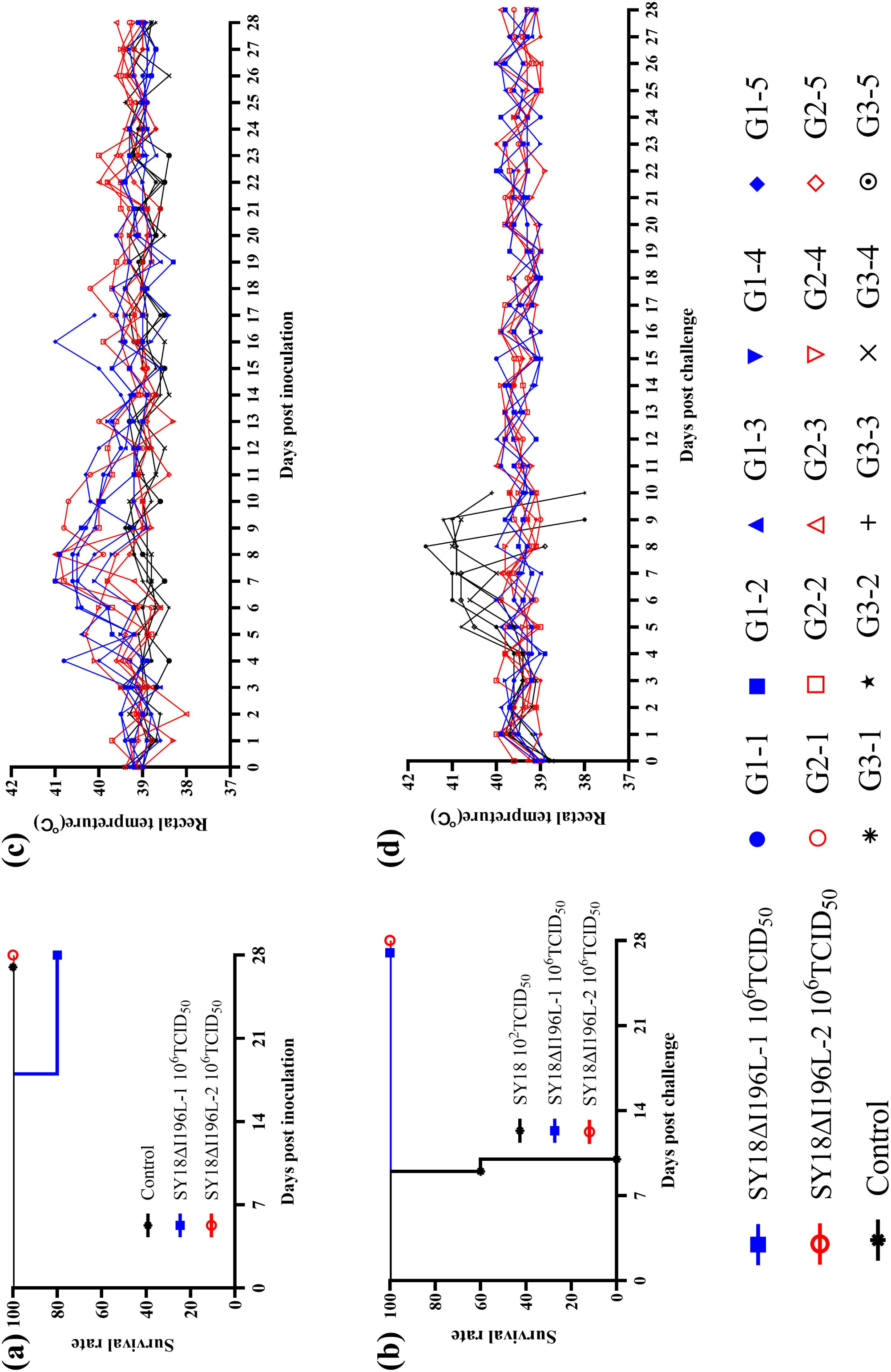
Survival rate and body temperature changes of pigs during immunization and challenge. (a)Survival rate of the pigs in three groups during immunization (inoculation with 10^6^ TCID_50_ SY18ΔI196L-1, 10^6^ TCID_50_ SY18ΔI196L-2 and saline in control). (b) Survival rate of the pigs after challenge (inoculation with 10^2^ TCID_50_ ASFV SY18). (c) The rectal temperatures of pigs after immunization, where the curves represent on rectal temperature of everey pig. (d) The rectal temperatures of pigs after challenge.

**Table 2.** Clinical performance and survival rate after challenge

| Groups | No. of survivors/total | No. of fever/total | Mean value for fever parameter (SD)Fever ( $\geq 40.5^{\circ}\text{C}$ ) | | |
| --- | --- | --- | --- | --- | --- |
| | | | Day of onset (days±SD) | Duration no. of days (days±SD) | Highest rectal temp ($^{\circ}\text{C}\pm$SD) |
| $10^6$ TCID<sub>50</sub> SY18ΔI196L-1 | 4/4 | 0/4 | —<sup>a</sup> | —<sup>a</sup> | —<sup>a</sup> |
| $10^6$ TCID<sub>50</sub> SY18ΔI196L-2 | 5/5 | 0/5 | —<sup>a</sup> | —<sup>a</sup> | —<sup>a</sup> |
| Mock | 0/5 | 5/5 | 5.8±0.75 | 3.2±0.4 | 41.12±0.27 |

a : The pigs did not have a fever.

The viremia was shown during the immunization and challenge periods in Fig. 6a and 6d. During immunization, due to the individual differences, the level of viremia varied significantly among individuals ranging from 10^4.73^ copies/mL to 10^7.32^ copies/mL on day 14, then steadily decreased. In particular, ASFV DNA could not be detected in the blood of pig G1-1 on day 21 and pig G2-5 on day 28. During challenge, pigs in G3 developed severe clinical symptom and high viremia ranging from 10^6.63^ copies/mL to 10^8.18^ copies/mL. After challenge, the viremia in G1 and G2 showed an overall decreasing trend, but individual changes were very interesting. Pig G1-1 had no detectable presence of virus in the blood throughout the 28 days of the challenge. During the challenge, virus levels in the blood of pigs G1-2, G1-4, G2-1 and G2-3 increased slightly and then decreased. virus levels in the blood of pigs G1-3, G2-2, and G2-4 continued to decrease, with no virus detected in the blood of pig G1-3 for 21 days of the challenge and no virus detected in the blood of pig G2-4 for 7 days of the challenge. No virus was present in the blood of pigs G2-5 on day 0 of the challenge (also day 28 of immunization), and then detected in the blood during the challenge (viral loads of 10^3.81^ copies/mL, 10^5.20^ copies/mL, and 10^4.25^ copies/mL in the blood on days 7, 14, and 21, respectively), but in the presence of virus was not detected in the blood on day 28.

**FIG 6.**
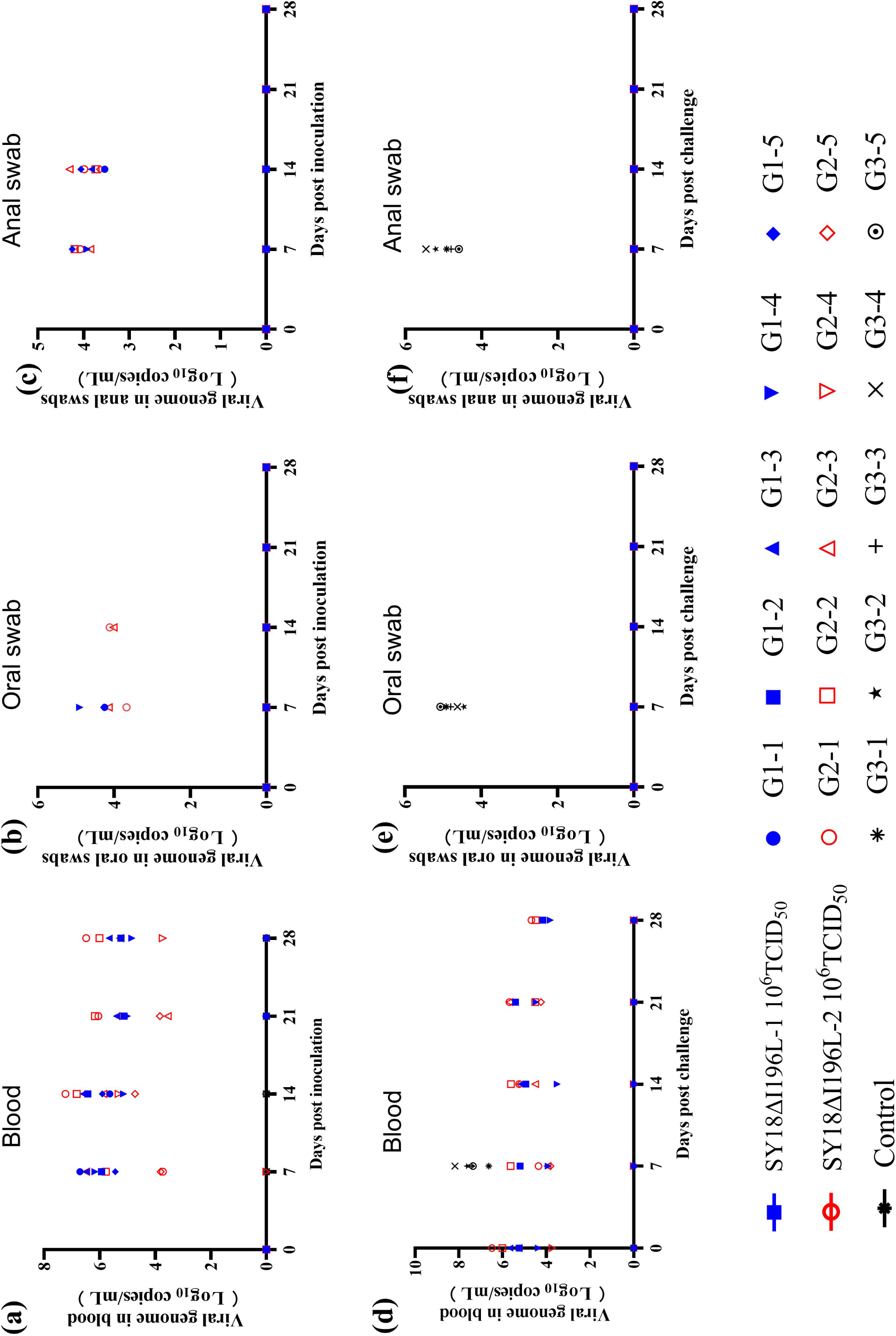
Detection of ASFV genomic DNA in blood, oral swabs and anal swabs. (a, d) Viral load of blood of each group during the immunization and challenge. (b, e) Viral load of oral swabs during the immunization and challenge. (c, f) Virus content in anal swabs during the immunization and the challenge. The y-values were indicated log10 copies/mL and the x-values were times of inoculation and challenge.

And we also examined the copies of viral DNA in oral (Fig. 6b and Fig. 6d) and fecal swabs (Fig. 6c and Fig. 6f) during the immunization and the challenge. ASFV DNA were detected in oral swabs and anal swabs on days 7 and 14 post immunization in occasional pigs, but ASFV DNA copies were low (10^3.53^-10^4.23^ copies/mL). No virus shedding was detected on days 21 and 28 post immunization. Among the pigs G1-2 and G2-4, virus shedding was never detected during the immunization period, which demonstrates that individual differences in the detoxification status of pigs against SY18ΔI196L virus can occur. No ASFV DNA were detected in pigs of groups 1 and 2, while the pigs of control group were detected in oral (10^4.45^-10^5.07^ copies/ mL) and anal swabs (0^4.60^-10^5.46^ copies/ mL) after the challenge.

All the dead pigs and control pigs were dissected and the level of ASFV DNA was detected in heart, liver, spleen, lung, kidney, submandibular lymph node, inguinal lymph node, thymus, bone marrow, colon and mesenteric lymph node with quantitative real-time PCR. The results were shown in Fig. 5. There was lower viral load in the tissues of pigs in group 1 and 2 compared to the challenged pigs. Furthermore, some tissues were negative.

### The level of p54 antibody in the animals infected with recombinant virus

The specific immune mechanism of ASFV-infected animals is still unclear, but specific antibody expression is the important indicator of immune response. Investigators found that the presence of ASFV-specific circulating antibodies may be associated with protective properties(23, 24). Therefore, we detected the level of p54 antibody in sera collected from the animals every 7 days post of infection and challenge. The results showed that the p54 antibody were negative on day 7 post immunization in group 1 and group 2, gradually increased from day 7 and all the pigs turned positive on day 14 (Fig. 8). There was little change before and after the challenge. The antibody levels in the serum of the control animals were always negative.

**FIG 7.**
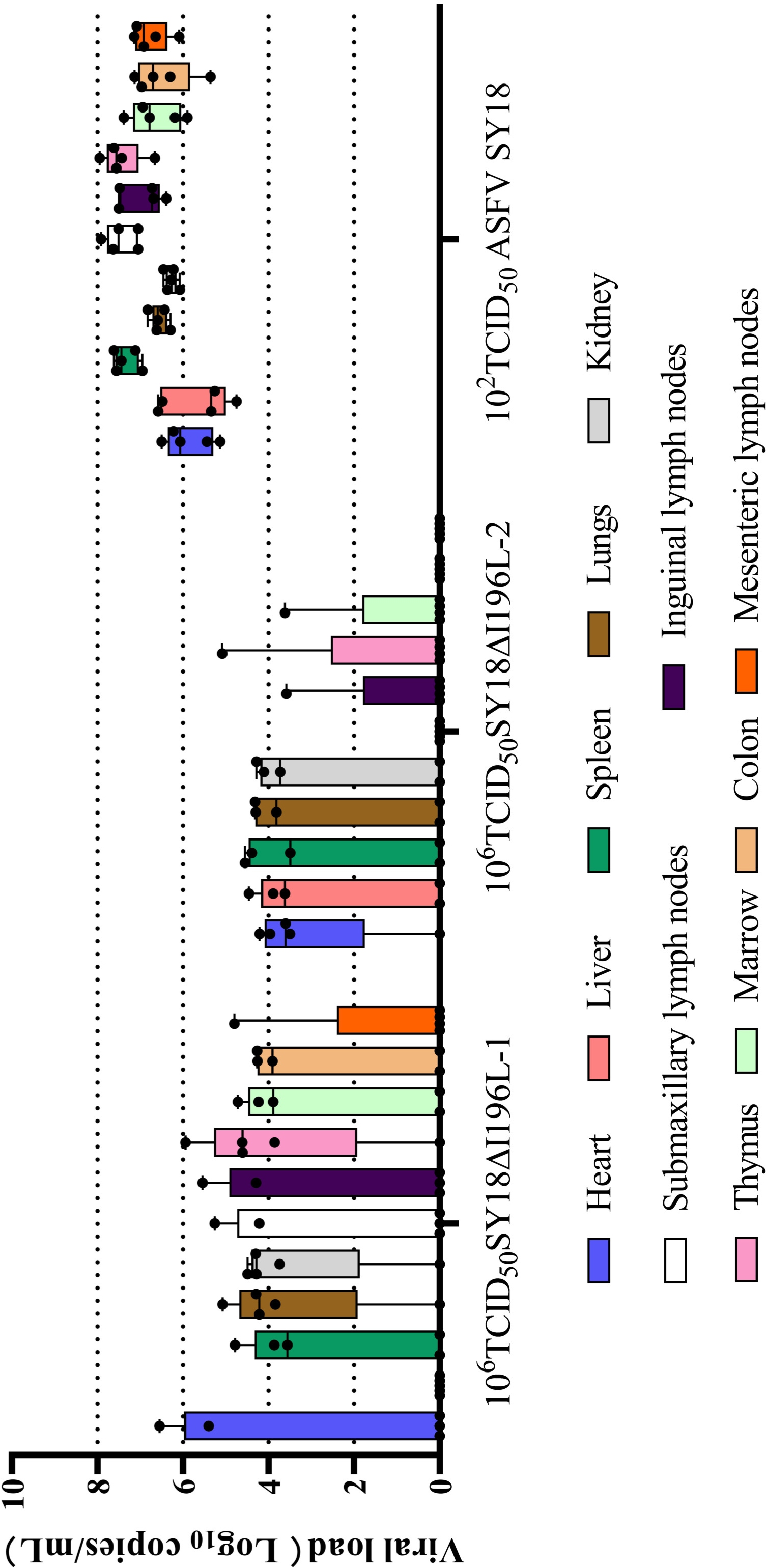
The results of viral load of tissues in three groups. The black dots represent viral load of the individual pigs. The values indicate log10 copies/mL. The upper edge of the box plot corresponds to the data that ranks in the 25th percentile when arranged in ascending order, and the lower edge corresponds to the data that ranks in the 75th percentile. The black horizontal line in the box plot represents the median.

**FIG 8.**
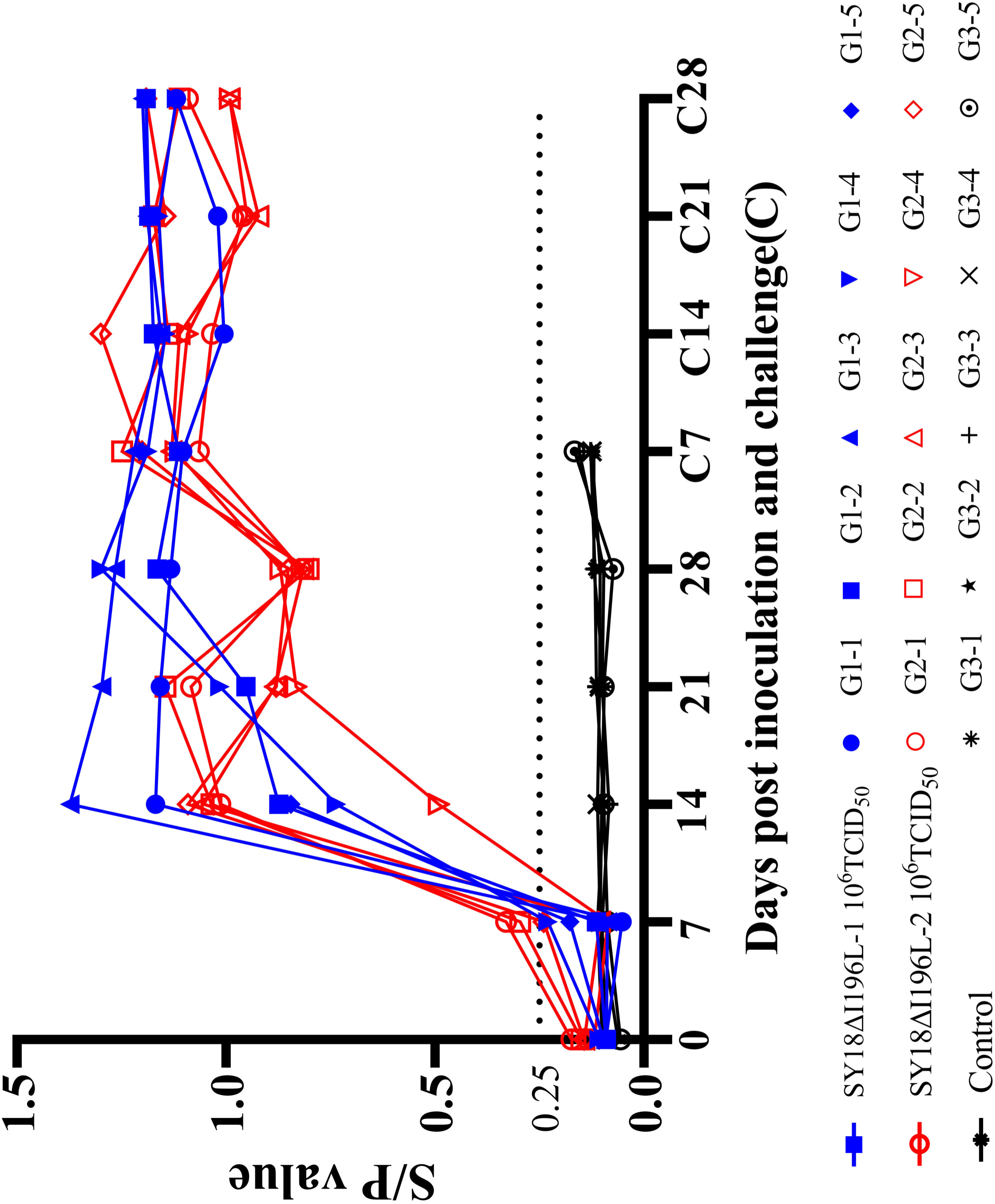
The level of antibody of ASFV p54 in the serum of pigs post the immunization and the challenge. Every symbol represented every pig and the symbol of same color represented the pigs in the same group. The blue lines represented the p54 antibodies of the pigs inoculated with 10^6^ TCID_50_ SY18 ΔI196L-1. The red lines represented the p54 antibodies of the pigs inoculated with 10^6^ TCID_50_ SY18 ΔI196L-2. And the black lines represented the p54 antibodies of the pigs in control group. The y-axis represented the ratio of S/P (sample OD450/positive control OD450) and the x-axis represented the days post inoculation and challenge.

## DISCUSSION

ASF seriously threatens global pork production and food security, especially in China. China not only is the biggest country of pig and pork production but also is the largest consumption market of pork in the world. The lack of a safe and effective commercial vaccine and the global pandemic of the disease stimulate the development of laboratorial researches on vaccines against ASF, including inactivated vaccines(25), live attenuated vaccines(19, 20, 26), virus vectored vaccines(27, 28) and subunit vaccines(29, 30). Among these, live attenuated vaccines provide better immunogenicity and more stable efficacy(31-33).

In this experiment, we identified a previously unknown functional ASFV gene, I196L, which is involved in virulence. The I196L is a late-expressed gene, which indicates that the gene may be little to do with ASFV DNA replication and closely related to viral assembly. We delete I196L with two different length, including SY18ΔI196L-1 deleted the I196L ORF except the shared 8 based pairs and SY18ΔI196L-2 deleted the whole ORF including the shared 8 based pairs. The two recombinant viruses had similar replication but all decreased comparing to the parental SY18 in vitro, which may mean I196L playing important function.

SY18ΔI196L-1 and SY18ΔI196L-2 have significant reduction in virulence. The pigs were inoculated the recombinant viruses with the high doses (10^6^ TCID_50_). All pigs were free of ASF-related symptoms except a transient mild fever of part of pigs between 1 and 3 days. In SY18ΔI196L-1 group, one pig died suddenly on the second day after blood collection. It was worth noting that the pig had a good condition before blood collection. We conjecture that the death of the pig was caused by the stress due to the blood collection of the experimental staff on the 14th day post immunization. We also found that pigs infected with African swine fever, whether strong strains or attenuated strains, are extremely sensitive to stress response and often end up sudden death. Thus, the dead pig is not counted as infected death. 1/5 pig in group 1 and 2/5 pigs in group 2 never developed high fever, the other pigs only had a transient fever after immunization and all the pigs were clinically normal. The result in group 1 inoculating SY18ΔI196-1 showed deleting I196L almost completely reduced the virulence of ASFV SY18.

The result in group 2 inoculating SY18ΔI196-2 is similar with group 1. The recombinant SY18ΔI196-2 simultaneously destroyed the genes of I196L and I177L. I177L gene has two kinds of annotation based on the location of initiation codon. The initial I177L ORF has only 201 bp encoding a 66 amino acid protein (being the 112th amino acid to the 177th amino acid position in the latest I177L ORF). Afterwards, the adjacent ASFV-G-ACD-0176 ORF was counted into the I177L genome and the latest I177L ORF became 534 bp encoding a 177 amino acid protein(11, 34, 35). The long ORF (534 bp) contain the short ORF (201 bp), the former is common. Here we assume that the short ORF of I177L is a key of the virulence of ASFV. There is report that deletion of the I177L gene about 112 nucleotides (located in short ORF) of 2007 Georgia isolate results in sterile immunity against the current epidemic eurasia strain and no fever happened. Here, SY18 ΔI196L-2 destroyed the long ORF of I177L gene and remained the short one, meanwhile, deleting the whole I196L gene. The results showed that more than half the pigs had a fever and there was also no significant difference in the effect of SY18ΔI196L-2 and SY18ΔI196L-1. The result demonstrated that I196L and short I177L ORF are the key genes of virulence.

Virus shedding and prolonged viremia are common during the observation of live attenuated ASFV vaccines, include HLJ/-18-7GD(36), ASFV-G-ΔA137R(16), SY18Δ I226R(20) and ASFV-GZΔI73R(21). We found that the deletion of the ASFV CD2v gene associated with the function of erythrocyte adsorption can significantly reduce the persistence of viremia. SY18ΔI196L-1 and SY18ΔI196L-2 induced a strong specific humoral immune response, though the antibodies do not neutralize ASFV, they play an important role in immunity. The pigs innoculated with SY18ΔI196L-1 and SY18ΔI196L-2 successfully resisted the lethal attack with normal body temperature and good clinical, meanwhile, viremia showed an overall downward trend and no detectable viral shedding in the oral cavity or feces. This result demonstrated that the two recombinant viruses had protective ability and safety.

An undesired mutation was detected in the SY18ΔI196L-2 genome resulting in a change in the 133rd amino acid of the E199L gene from Gly to Arg. Although it has been demonstrated that pE199L is a protein that affects autophagy and is relevant for membrane fusion and core penetration(37, 38). However, we believe that this mutation of one amino acid in E199L is not a critical factor affecting the virulence of SY18ΔI196L-2. In summary, the research demonstrates that I196L in ASFV genome is a key gene on virulence. The deletion of I196L in ASFV SY18 maintains good immunogenicity while significantly reducing virulence. It is a novel and effective live attenuated vaccine (LAVs).

## MATERIALS AND METHODS

### Cells and viruses

Primary alveolar macrophages (PAMs) were prepared from healthy porcine lungs as described before(20). PAMs were cultured in RPMI 1640 medium added 5% fetal bovine serum (FBS) and *penicillin-streptomycin* (100 U/ml of *penicillin* and 100 μg/ml of *streptomycin*).

The ASFV SY18 strain (GenBank number: MH766894) is virulent field isolate from Changchun Veterinary Research Institute, Chinese Academy of Agricultural Sciences.

### Viral content

The PAMs were seeded into 96-well plates in advance. Three ASFV strains were diluted in a 10-fold gradient and 100 μL dilution of per gradient was added to 8 wells of 96-well plates. The PAMs were cultured for 4 days and then stained by anti-p30 monoclonal antibody labeled with fluorescein isothiocyanate (FITC) (prepared by our laboratory). The viral titers were calculated using the Reed-Muench method.

### Sequence analysis

The I196L in different ASFV strains were collected in the NCBI database and translated into amino acid. The MAFFT website and the Jalview software were used to compare and evaluate the conservativeness of the I196L amino acid residues.

### I196L expression

PAMs were seeded (1 × 10^6^ cells/well) in 12-well plates after resuscitation infected with ASFV SY18 at 3 MOI and the negative controls. The infected cells were collected at 1, 4, 8, 12, 16 and 24 h. Three replicates were set up for all groups. RNA extraction was performed using FastPure Cell/Tissue Total RNA Isolation Kit V2 (Vazyme, Nanjing, China) and cDNA synthesis was performed using HiScript III RT SuperMix for qPCR (+gDNA wiper) (Vazyme). The real-time fluorescent quantitative PCR basing SYBR Green Dye targeted GAPDH, p30 (CP204L), p72 (B646L) and I196L. The primers of part of gene were performed as previously described(8) and the primer of I196L gene was designed by Primer Premier 6 software, which were shown in Table 1.

**Table 1.**
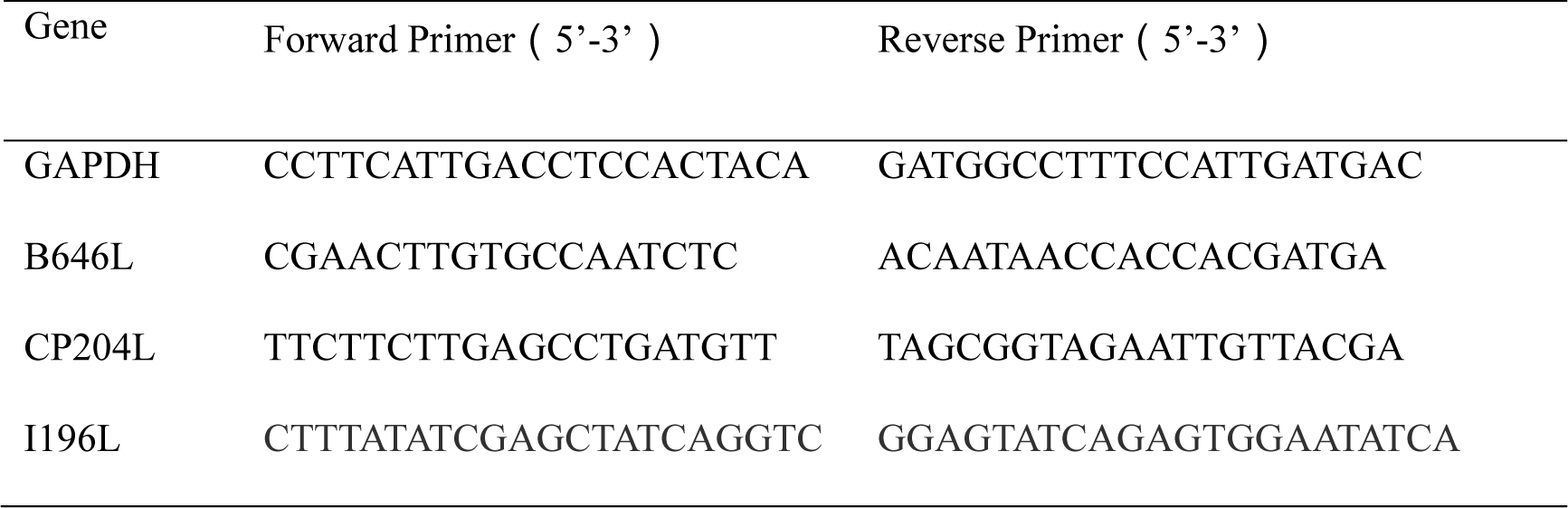
Primer information

### Construction and purification of SY18ΔI196L-1 and SY18ΔI196L-2

Recombinant transfer vectors (p72eGFPΔI196L-1, p72eGFPΔI196L-2) were constructed containing upstream and downstream fragments of the target deletion I196L gene and a reporter gene cassette containing the promoters of the eGFP and ASFV p72 late-stage genes. The I196L gene in SY18 strain contains a total of 609 bp containing 202 amino acids. p72eGFPΔI196L-1 contains a flanking region of amino acids 1 to 200 of the I196L gene and p72eGFPΔI196L-2 contains a flanking region of amino acids 1 to 202 of the I196L gene (see Fig. 2a and 2b for detailed construction). PAMs were added in 12-well plates and incubated for 12 h. After infection with ASFV SY18 at 1 MOI for 4 h, the recombinant plasmids were transfected and the appearance of fluorescent cells could be observed after 24 h. Fluorescent cells were collected and purified by a limited dilution method. The purification of SY18ΔI196L-1 and SY18ΔI196L-2 strains was identified by polymerase chain reaction. Detection of primer 1: F, TCACTGTAGTAGGTCTATAAGG; R, CTTCATCCGACGATGAGTCC. A 549bp fragment could be amplified if the parental strain SY18 was present.

### Next-generation sequencing of SY18ΔI196L-1 and SY18ΔI196L-2 genomes

SY18ΔI196L-1 and SY18ΔI196L-2 were infected with PAMs and cell cultures were collected after 5 days. The viral genome was extracted using the somatic virus DNA/RNA miniprep kit (Axygen, USA).The full length of the genome was determined by Next-generation sequencing (NGSs) (Novogene, Tianjin, China).

### Growth characteristics of SY18ΔI196L and SY18ΔI196L-2

To evaluate the growth characteristics in vitro, low dose of two recombinant viruses were used to infect PAMs, the parental virus was used as reference. The virus content was determined at different time. PAMs (10^6^ cells/well) were seeded into 12-well plates and incubated for 12 h. Cells were infected with ASFV SY18 and SY18ΔI196L at 0.01 MOI and cell cultures were collected at 1, 6, 12, 24, 48 h, 72 h, 96 h and 120 h. Three replicates were set up for all groups. After repeated freeze-thaws, virus titers were measured at each time point.

### SY18ΔI196L and SY18ΔI196L-2 genome sequencing

SY18ΔI196L was infected with PAMs and cell cultures were collected after 5 days. The viral genome was extracted using the somatic virus DNA/RNA miniprep kit (Axygen, USA). The full length of the genome was determined by second generation sequencing technology.

### Electron microscopy of recombinant viruses

PAMs were infected with SY18ΔI196L-1, SY18ΔI196L-2 and ASFV SY18 at 5 MOI and the cells were collected into 1.5mL conical centrifuge tubes after 48 hpi and centrifuged at 5,000 × g for 10 min at 4 °C. After dumping the supernatant, the cell precipitation was fixed in 2.5% glutaraldehyde solution at 4℃ overnight and were further treated to prepare for ultrathin sections and observed by electron microscopy. Briefly, the samples were rinsed with phosphoric acid buffer of 0.1M and pH7.0, fixed with 1% osmic acid solution for 1 h, rinsed with double steaming water twice for 10 min per time, dehydrated with ethanol solution of gradient concentration (including 50%, 70%, 80%, 90% and 100%) at 4℃, soaked in pure acetone at room temperature for 20 min and then in the mixture of embedding agent and pure acetone (1:1 volume ratio) at room temperature for 30 min, permeated in pure embedding agent overnight and polymerized at 35℃ for 12h, 45℃ for 12h, and 60℃ for 48h. Lastly, the embedded samples were modified and sliced to obtain sections of 70∼90 nm, which were stained with uranium dioxyacetate solution and lead citrate solution for 10 min and 20 min respectively. The sections were observed on electron microscopy.

### Animal experiment

Fifteen Landrace pigs weighting about 15-17 kg were randomly divided into three groups to assessed the virulence of SY18 ΔI196L-1 and SY18ΔI196L-2. All experimental procedures involving ASFV were performed in biosafety level 3 laboratory at Changchun Veterinary Research Institute. The animal experiment was conducted according to the standard procedures approved by the Animal Welfare and Ethics Committee of the Changchun Veterinary Research Institute (review ID: IACUC of AMMS-11-2020-018). The pigs in two groups were intramuscularly inoculated with 10^6^ TCID_50_ of SY18 ΔI196L-1 (group 1, G1), and 10^6^ TCID_50_ of SY18 ΔI196L-2 (group 2, G2), respectively and the pigs in last group were intramuscularly inoculated with an equal amount of saline (group 3, G3). After 28 days post infection (dpi), all animals survived were intramuscularly challenged with 10^2^ TCID_50_ of ASFV SY18 and continued to be observed for 28 days post challenge (dpc). The clinical signs and rectal temperature were recorded daily. Blood, oral swabs, and anal swabs were collected every 7 days. The pigs were euthanized with pentobarbital while becoming moribund and ending the observation and the tissues were collected for ASFV detection and fixed by 4% paraformaldehyde for pathological analysis. The copies of p72 gene in blood, oral swabs, anal swabs and tissues were measured by qPCR recommended by the National Standard of the People’s Republic of China (GB/T 18648-2020) (https://openstd.samr.gov.cn/bzgk/gb/index).

### Detection of p54 antibodies in ASFV

The measurement of the p54 antibody was performed using sandwich ELISA based on p54 protein, one of structural protein of ASFV, prepared by our laboratory. The p54 antibodies in each serum sample was detected as previously described in the procedure(29). Absorbance of the samples was measured at OD450. And the sample was considered a positive sample when the ratio of S/P (sample OD450/positive control OD450) exceeded 0.25.

## DECLARATIONS

The study was conducted according to the standard procedures approved by The Animal Welfare and Ethics Committee of Changchun Veterinary Research Institute and the animal biosafety level 3 (ABSL-3) lab. All opinions expressed in this paper are those of the authors and do not necessarily reflect the policies and views of the Ministry of Agriculture and Rural Affairs of China.

## ACKNOWLEDGMENTS

We especially thank Prof. Shoufeng Zhang, Prof. Jinxia Zhang, Ms. Fei Zhang and Ms. Lijuan Mi for their kind assistance. This work was funded by the National Key Research and Development Program of China (Grant No. 2021YFD1800100 and 2021YFD1801204)

The authors declare no competing interests.

## AUTHOR CONTRIBUTIONS

Conceived and designed the experiments: RLH and TC. Performed the experiments: FJQ ZRN LN YJJ YHX ZYY ZXT KJN WY LQX and QY. Analyzed the data: FJQ ZRN LN and LM. Contributed reagents/materials/analysis tools: MFM KJN WY LQX and QY. Wrote the paper: FJQ and ZYY.

